## Supplementary Figures for "Prediction of representative phenotypes using Multi-Attribute Subset Selection"

### Supplementary Tables

**Supplementary Table 1:**

DATASET 1 strains with additional metadata.

**Supplementary Table 2:**

Digitized values of table in Chapter 6 of (Barnett, Payne, and Yarrow 1990) resulting 590 yeast and 92 phenotypes (raw data for DATASET 3).

**Supplementary Table 3:**

Selection of attributes of Supplementary Table 2 used for MASS application.

### Supplemental Figures

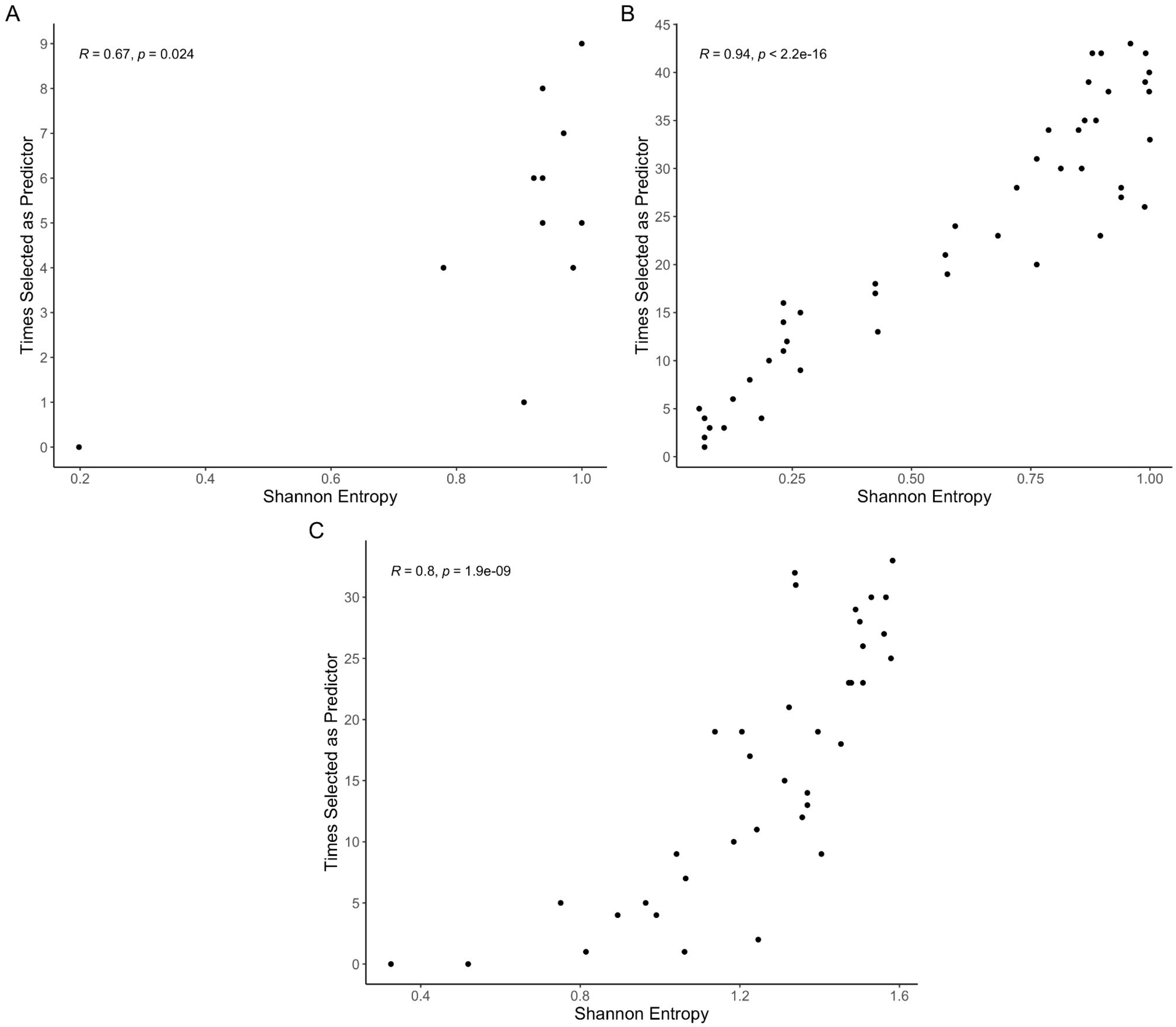


**Supplementary Figure S1. The variable predictor likelihood versus its information content**. The number of times a variable was selected as a predictor as a function of Shannon entropy for the marine bacterial growth DATASET 1 (**A**), the bacterial fermentation DATASET 2 (**B**), and the yeast phenotype DATASET 3 (**C**). The plots also contain the Pearson correlation coefficients for comparison.

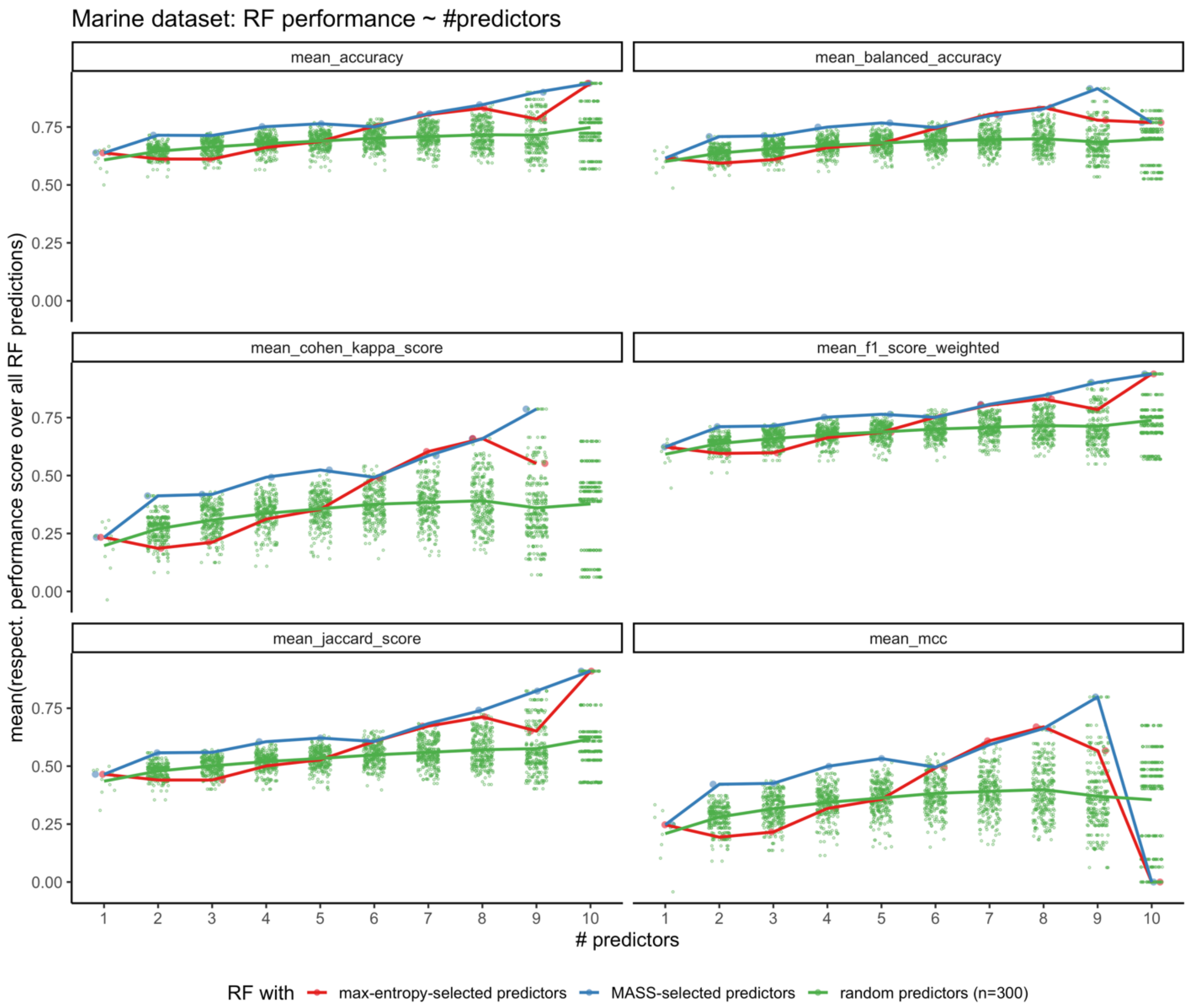


**Supplementary Figure S2. Various performance measures of random forest classifiers for DATASET 1**. Random forest (RF) classifiers were trained using environmental conditions selected by MASS (blue), maximal Shannon entropy (red), or random conditions as predictors. The classifiers were then evaluated using (from top-left to bottom-right panel) Accuracy, Balanced Accuracy, Cohen's Kappa score, weighted F1 score, Jaccard score and Matthews correlation coefficient (MCC). The MCC plot was used as Figure 2D.

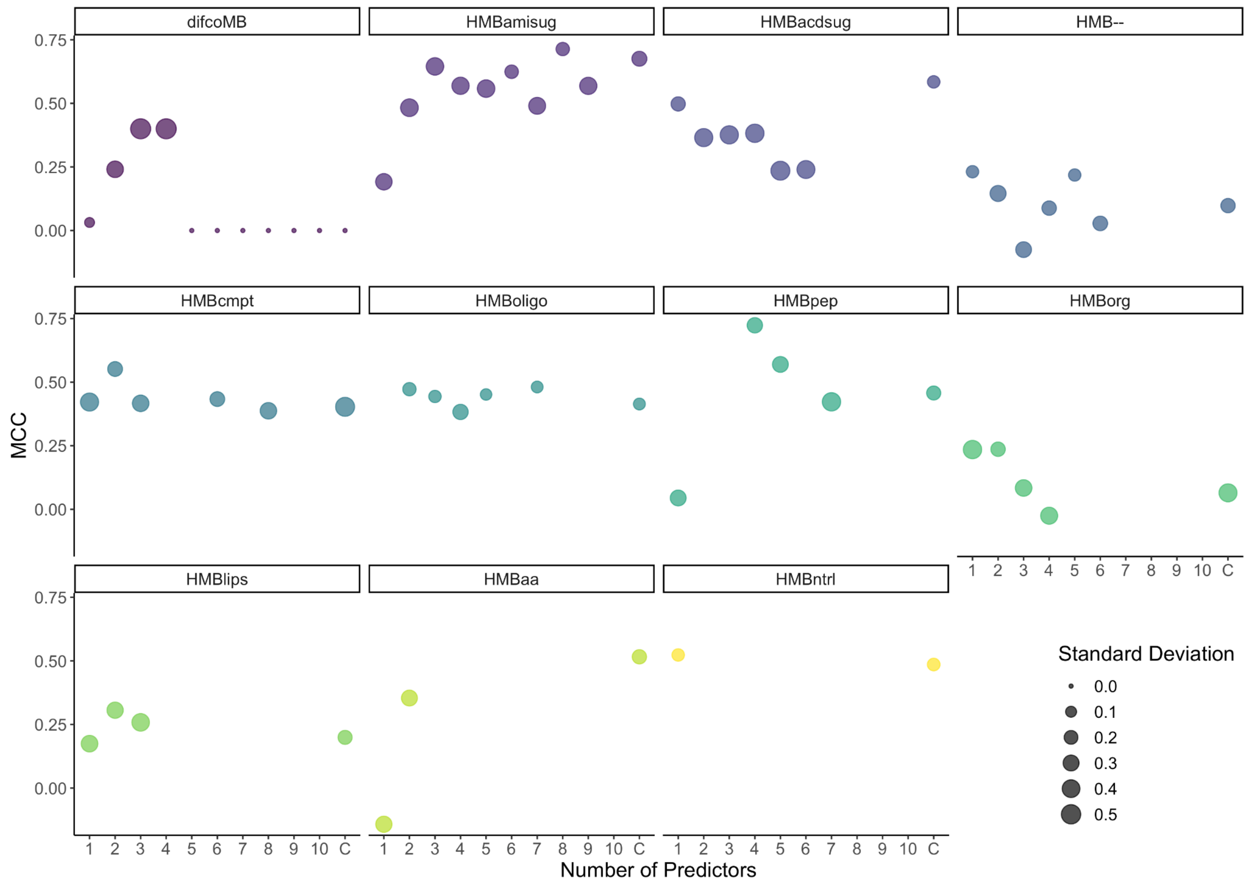


**Supplementary Figure S3. DATASET 1 performance measures of random forest classifiers for individual responses**. For each possible response condition, the Matthews correlation coefficient (MCC) of each random forest model for each number of predictors, *p*, is depicted. Dot size represents the standard deviation of performance scores for values obtained via 5-fold cross-validation; column C is the RF control using all other conditions as predictors to predict a response condition. If no dot is displayed for a condition, the condition was selected as a predictor. Media conditions are ordered based on their likelihood to be selected as response by MASS and named as follows: Difco Marine Broth (difcoMB), eight engineered media with single classes of carbon sources (HMBpep = peptides; HMBaa = amino acids; HMBlips = lipids; HMBoligo = oligosaccharides; HMBorg = organic acids; HMBntrl = neutral sugars; HMBamisug = amino sugars; HMBacdsug = acidic sugars), a defined medium containing all 8 carbon classes (HMBcmpt), and a medium with no added carbon sources (HMB–).

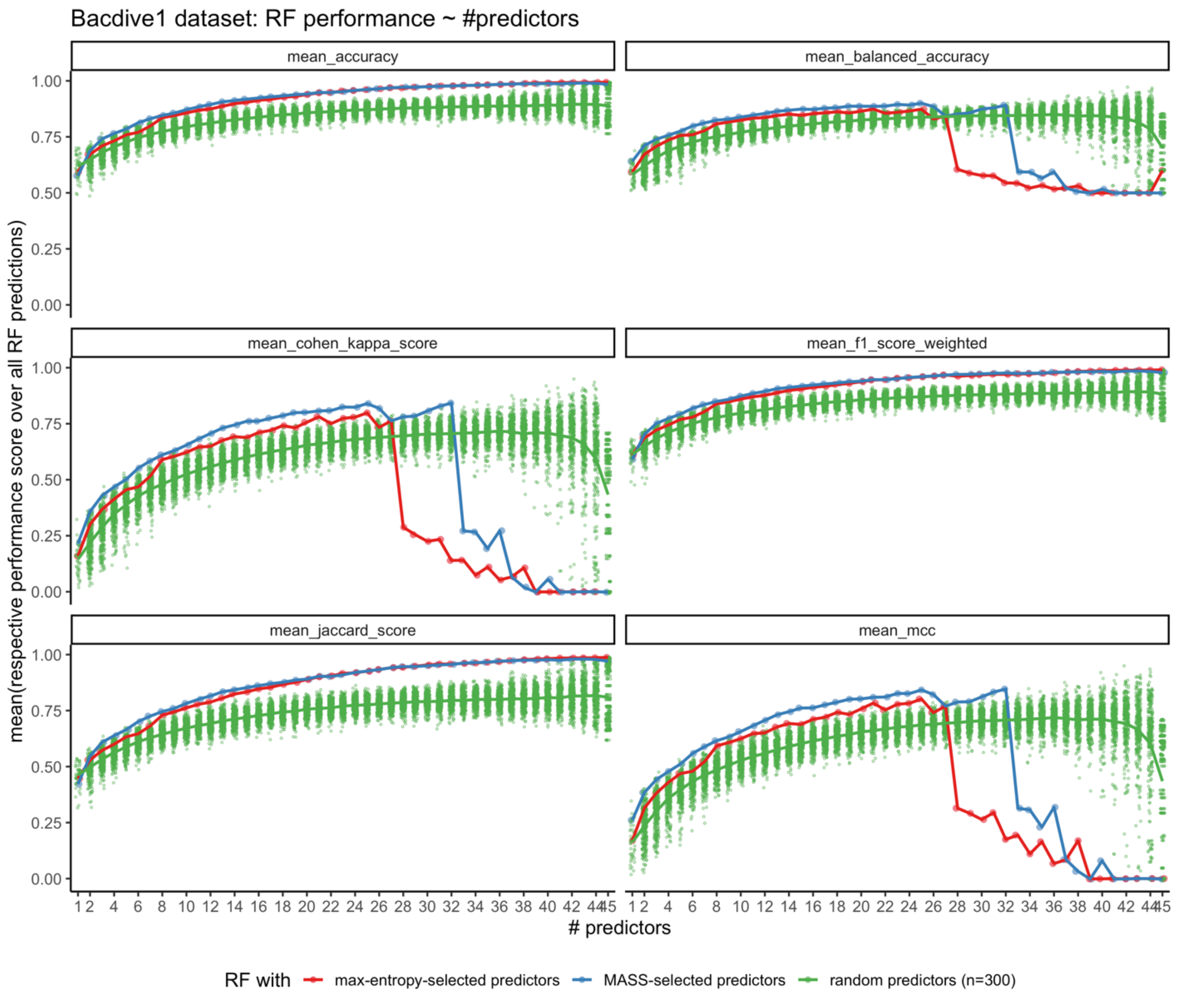


**Supplementary Figure S4. Various performance measures of random forest classifiers for DATASET 2**. Random forest (RF) classifiers were trained using environmental conditions selected by MASS (blue), maximal Shannon entropy (red), or random conditions as predictors. The classifiers were then evaluated using (from top-left to bottom-right panel) Accuracy, Balanced Accuracy, Cohen's Kappa score, weighted F1 score, Jaccard score and Matthews correlation coefficient (MCC). The MCC plot was used as Figure 3D.

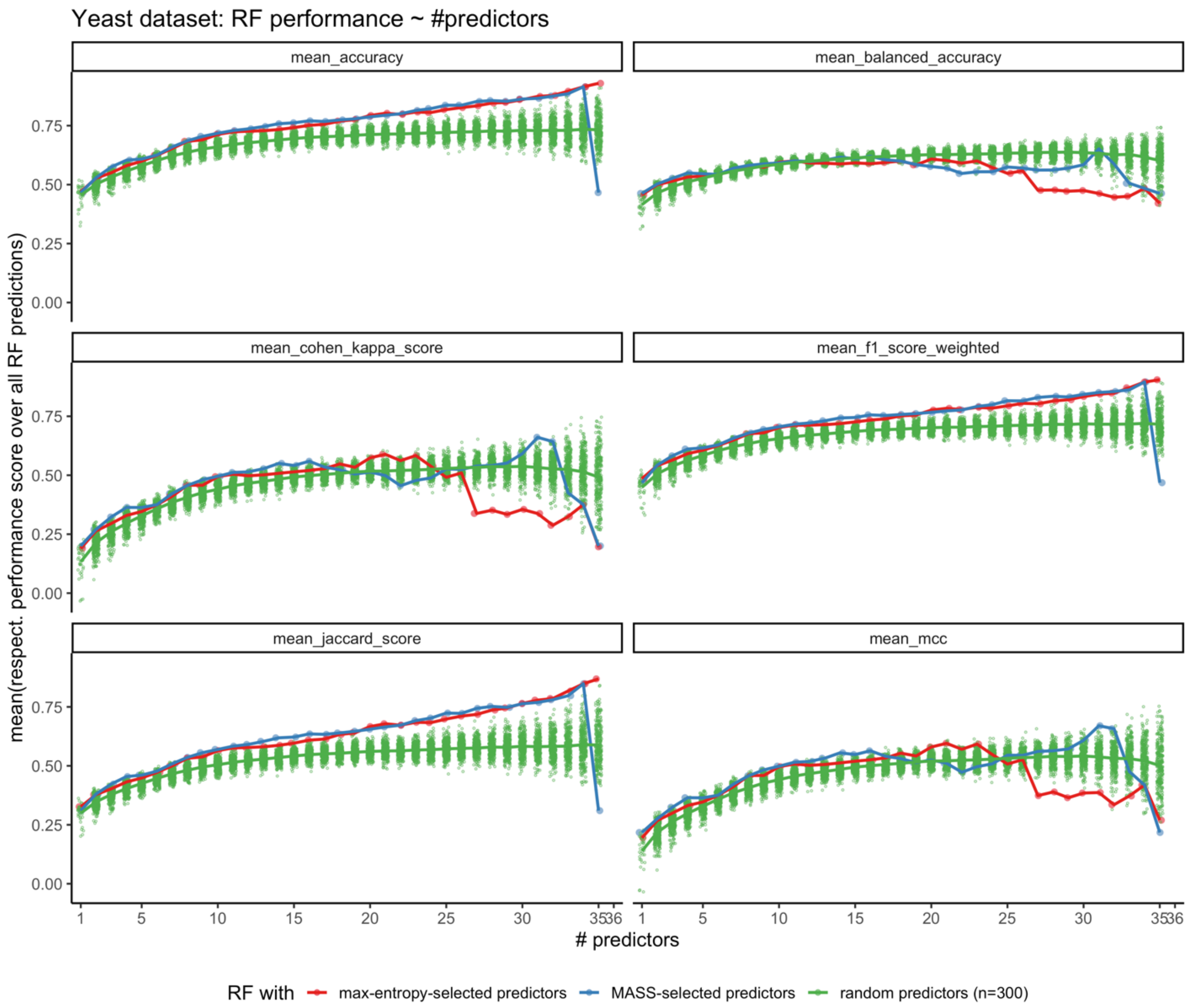


**Supplementary Figure S5. Various performance measures of random forest classifiers for DATASET 3.** Random forest (RF) classifiers were trained using environmental conditions selected by MASS (blue), maximal Shannon entropy (red), or random conditions as predictors. The classifiers were then evaluated using (from top-left to bottom-right panel) Accuracy, Balanced Accuracy, Cohen's Kappa score, weighted F1 score, Jaccard score and Matthews correlation coefficient (MCC). The MCC plot was used as Figure 4D.
